# Quality Assessment of Different Brands of Diclofenac Tablets Marketed in Sagamu Community

**DOI:** 10.1101/2025.07.17.665276

**Authors:** L.S. Kasim, M.V. Badejo, T.O. Paramole, J.O. Daodu, K. O. Olufolabo, O. L. Okunye, T.A. Banjo

**Affiliations:** Department of Pharmaceutical and Medicinal Chemistry, Faculty of Pharmacy, Olabisi Onabanjo University, Sagamu, Ogun State; Department of Pharmacognosy, Faculty of Pharmacy, Olabisi Onabanjo University, Sagamu, Ogun State; Department of Pharmaceutical Microbiology, Faculty of Pharmacy, Olabisi Onabanjo University, Sagamu, Ogun State; Department of Medical Microbiology and Parasitology, Faculty of Basic Clinical Sciences, Olabisi Onabanjo University, Sagamu, Ogun State

**Keywords:** analgesic activity, chemical equivalence, diclofenac potassium, quality control

## Abstract

The increasing prevalence of counterfeit and substandard pharmaceutical products poses a significant global health challenge. Given the multitude of diclofenac brands available in the market, ensuring their quality, safety, and efficacy through rigorous quality control testing is essential to safeguard public health and maintain therapeutic reliability. This study aimed to evaluate the *in-vitro* chemical equivalence and *in-vivo* analgesic efficacy of diclofenac potassium tablets (50 mg) across five different brands. Comprehensive quality control tests, including weight variation, friability, hardness, disintegration, dissolution, and content purity analysis, were performed in accordance with the British Pharmacopoeia standards. The *in-vivo* analgesic activity was assessed using the hot plate method. All five brands (A, B, C, D, and E) achieved a 100% pass rate in weight variation, friability, hardness, and disintegration tests. However, Brands A, C, D, and E failed the dissolution test, with drug release percentages of 51.2%, 52.21%, 30.53%, and 46.17% at 45 minutes, respectively, while Brand B met the official standard with a drug release of 70.44%. Content purity analysis revealed that Brands A, B, C, and E conformed to the pharmacopoeia standards, with percentage purities of 97%, 95.5%, 100%, and 98.5%, respectively, while Brand D failed with a content purity of 75%. Despite these variations, all brands demonstrated significant analgesic activity in vivo, indicating potential therapeutic efficacy. The study highlights the necessity for continuous post-market surveillance and strict compliance with quality standards to ensure the safety, efficacy, and reliability of pharmaceutical products.

## Introduction

Diclofenac, a derivative of phenylacetic acid, is a widely used non-steroidal anti-inflammatory drug (NSAID) available in various salt forms, including diclofenac sodium and diclofenac potassium [1]. Diclofenac sodium was first introduced in Japan in 1974 as a slow-release tablet aimed at managing chronic pain by ensuring sustained drug release over an extended period [2]. Conversely, diclofenac potassium was developed to facilitate rapid absorption, making it particularly effective for conditions requiring swift pain relief [1]. In addition to its analgesic properties, diclofenac also exhibits notable anti-inflammatory and antipyretic activities [1].

The pharmacological effects of diclofenac are mediated through the inhibition of cyclooxygenase-1 (COX-1) and cyclooxygenase-2 (COX-2), key enzymes involved in the biosynthesis of prostaglandin-E2 (PGE2), prostacyclins, and thromboxanes, which play pivotal roles in inflammatory and nociceptive pathways [3,4]. Furthermore, diclofenac modulates the lipoxygenase pathway, enhancing its anti-inflammatory effects, although it does so without directly inhibiting the lipoxygenase enzyme [3].

Diclofenac is widely accessible with or without a prescription and is globally recognized as the most prescribed NSAID, commanding the largest market share among other NSAIDs [5]. In Nigeria, diclofenac is commonly prescribed in hospital settings. A retrospective study conducted at Lagos University Teaching Hospital (LUTH) revealed that diclofenac potassium was the second most prescribed NSAID in outpatient clinics, following aspirin [6]. Similarly, in the geriatric outpatient clinic of the University College Hospital (UCH) Ibadan, diclofenac ranked second to paracetamol in prescription frequency [7]. Furthermore, a study evaluating the use of potentially inappropriate medications (PIMs) among older patients at UCH identified diclofenac as the most frequently used PIM, accounting for 51.3% of all recorded PIMs [8].

Numerous brands of diclofenac potassium tablets are available in the Nigerian market, with some locally manufactured and others imported. However, the prevalence of counterfeit and substandard drugs underscores the importance of stringent quality control measures for these products. Studies evaluating the quality of diclofenac tablets in Nigeria have reported concerning findings. For instance, Ayorinde et al. (2012) revealed that 60% of tested diclofenac sodium brands failed the dissolution test which is a measure of drug release and bioavailability [9] . Similarly, Adeyemi et al. (2017) reported that only 13.3% of the sampled diclofenac brands passed the chemical equivalence test, assessed using high-performance liquid chromatography (HPLC) to determine chemical purity [10].

Another study found that 42.86% of the tested diclofenac samples failed to comply with the British Pharmacopoeia specifications regarding the percentage composition of diclofenac [11]. This observation is consistent with the findings of Abdullahi and colleagues, who reported the presence of substandard diclofenac tablets among the analysed samples [12]. Such evidence highlights the widespread of poor-quality diclofenac tablet formulations and underscores the need for continuous and rigorous quality control checks on diclofenac products marketed in Nigeria. Such measures are critical for ensuring consumer safety, promoting prescriber confidence, and upholding the integrity of regulatory authorities responsible for maintaining pharmaceutical standards. This study therefore focused on the quality assessment of five different brands of diclofenac potassium tablets marketed in Sagamu, Ogun State, Nigeria.

## Materials and Methods

### Chemical reagents/Instruments

Diclofenac Potassium (Sigma-Aldrich, U.S.A), High Performance Liquid Chromatography (HPLC Waters, U.S.A.), friability tester (DBK, India), hardness tester, dissolution test apparatus (Copley scientific, U.K.), disintegration tester (Copley scientific, U.K.) UV-Vis Spectrophotometer (Thermo Fisher Scientific, U.S.A.). All other reagents used for the study were of analytical and HPLC grade.

### Collection of samples

Five (5) different brands of diclofenac potassium (50 mg) uncoated tablets were purchased in June 2024 from different pharmacy outlets in Sagamu, Ogun state, Nigeria. Information regarding the manufacturing date, expiry date, batch number, country of manufacture, and the National Agency for Food and Drug Administration and Control (NAFDAC) registration number were documented. The samples were assigned codes A – E.

### Uniformity of weight test

Twenty (20) tablets of each brand of diclofenac potassium were weighed individually using a digital weighing balance (Mettler Toledo, Switzerland). The average weight and the percentage weight variation for each brand was calculated using the formular below [13]

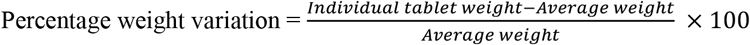

### Friability test

A total of ten (10) tablets were randomly selected from each brand and weighed to determine their initial weight. The tablets were then subjected to abrasion and shock in a friabilator (DBK, India) operating at 25 revolutions per minute (rpm) for a duration of 4 minutes. During each revolution, the tablets were dropped from a height of six inches. After the procedure, the tablets were removed from the friabilator, de-dusted, and reweighed to determine their post-friabilation weight [13]. The percentage weight loss, which serves as an indicator of tablet friability, was calculated using the formula:

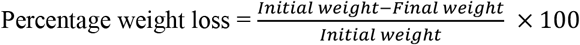

### Hardness test

The hardness of ten tablets from each brand of diclofenac potassium (50 mg) was evaluated using a tablet hardness tester. Each tablet was individually placed between the two jaws of the tester, and a force was applied until the tablet fractured. The hardness of each tablet was measured as the amount of force required to crush the tablet, recorded in newton [13].

### Disintegration test

Six tablets from each brand of diclofenac potassium (50mg) were subjected to disintegration test using the disintegration test apparatus made of transparent polyvinyl basket, which contained six tubes of uniform diameter. Each tube had a stainless-steel wire mesh with uniform mesh size where the tablets were placed for testing. Small metal discs were used to ensure the complete immersion of the tablets in the fluid medium (distilled water) maintained at a temperature of 37°C. The entire basket-rack assembly was mounted on a reciprocating motor positioned at the apex of the assembly. The motor facilitated the vertical movement of the basket at approximately 28 to 32 revolutions per minute (rpm) within the fluid medium. The disintegration time for each tablet was recorded, with disintegration being defined as the point at which the tablet completely broke down and no residue remained in the basket [14].

### Dissolution test

The dissolution test for various brands of diclofenac potassium tablets (50 mg) was conducted using the United States Pharmacopoeia (USP) Dissolution Apparatus. A phosphate buffer solution at pH 6.8 served as the dissolution medium, with a total volume of 900 mL maintained at a temperature of 37 ± 0.5°C to simulate physiological conditions. Each tablet was placed in a rotary basket, which was submerged in the dissolution medium and rotated at a speed of 50 rpm. At intervals, 5 mL aliquot of the dissolution medium was withdrawn and filtered to remove particulate matter. The amount of diclofenac potassium dissolved in the withdrawn samples was quantified using a UV-Vis spectrophotometer at a wavelength of 276 nm. To maintain a constant volume, 5 mL of fresh phosphate buffer (pH 6.8) was immediately added to the vessel after each sampling. A calibration curve for standard diclofenac potassium was plotted to establish the relationship between UV absorbance and known concentrations of the drug. This curve was used to calculate the concentration of diclofenac potassium dissolved at time intervals of 5, 10, 20, 30, 45, and 60 minutes.

### Quantitative analysis of diclofenac potassium brands using High Performance Liquid Chromatography (HPLC)

HPLC analysis was conducted following standard procedures [13]. The analysis utilized a Waters 265 HPLC system equipped with a 4.6 × 150 mm, 5 µm C18 column as the stationary phase. The mobile phase comprised acetonitrile and water in a ratio of 20:80, delivered by a quaternary pump at a flow rate of 1 mL/min. The system was operated at ambient temperature with UV detection at a wavelength of 280 nm and an injection volume of 20 µL. After setting and stabilizing the chromatographic conditions to obtain a stable baseline, a mixed standard solution of pure diclofenac potassium was prepared in the mobile phase and filtered. The solution was injected via a manual injector, and the chromatograms were recorded. Standard chromatograms of diclofenac potassium were generated for concentrations of 10, 40, 80, 120, 160, 200, 240, 280, and 300 µg/mL.

For the analysis of diclofenac potassium (50 mg) brands, 20 tablets of each brand were weighed and finely powdered. A quantity equivalent to 20 mg of diclofenac potassium was transferred into a 100 mL volumetric flask. Approximately 60 mL of distilled water was added, and the mixture was sonicated for 45 minutes. After cooling to room temperature, the solution was diluted to the 100 mL mark. The resulting solution was filtered through a 0.45 µm filter using a syringe and transferred into HPLC vials. A 20 µL aliquot of the final solution, corresponding to a concentration of 0.2 mg/mL, was injected into the HPLC system for analysis.

### Analgesic activity of different brands of diclofenac potassium (50 mg) on experimental rats

Thirty-five male albino rats weighing 118–123g were obtained from the Central Animal House, Faculty of Pharmacy, Olabisi Onabanjo University, Sagamu, Ogun State, Nigeria, where the experiment was also conducted. The animals were housed in clean, well-ventilated cages under standard environmental conditions and were acclimatized for two weeks prior to the commencement of the study. During this period, they were fed pelletized feed (Vital Feeds Ltd.) and provided with unrestricted access to water ad libitum.

The analgesic activity of various brands of diclofenac potassium was evaluated using the hot plate method. The animals were randomly divided into seven groups. Experimental Groups I to V received 1 mg/mL intraperitoneal injections of the respective diclofenac brands. Group VI, serving as the negative control, received an equivalent volume of distilled water, while Group VII, designated as the positive control, was treated with the standard reference formulation of diclofenac potassium.

Each rat was placed on a hot plate maintained at 55°C within a restrainer, and the reaction time (in seconds) or latency period was recorded. This was defined as the time taken for the rats to respond to the thermal stimulus by licking their paws or jumping. The reaction time was recorded before (0 min) and at 15, 30, 45, and 60 minutes after the administration of the diclofenac treatments. To ensure the safety of the animals, the maximum reaction time was capped at 45 seconds to prevent tissue injury [15].

### Statistical analysis

All data analyses were performed using GraphPad Prism software (version 8.0.1). An unpaired t-test was employed to compare the treatment groups with the control groups, with statistical significance set at a threshold of *p* < 0.05.

## Results

## Discussion

Continual post-production screening of pharmaceutical products is essential to ensure their quality, safety, and efficacy throughout their shelf life. In this study, five brands of diclofenac potassium tablets were procured from pharmacy outlets in Sagamu, Ogun State, Nigeria. Among these, 60% were manufactured in India, 20% in China, and 20% in Nigeria. At the point of purchase, all brands exhibited satisfactory aesthetic properties, good physical integrity, with no observable signs of dustiness, cracks, or breakages (Table 1). Qualitative and quantitative pharmacopoeia quality control tests were performed on the various brands to assess their compliance with established standards.

**Table 1:** Details of different brands of diclofenac potassium (50 mg) tablets used in the study.

| Brand code | Batch number | NAFDAC registration number | Manufacture date | Expiry date | Place of manufacture |
| --- | --- | --- | --- | --- | --- |
| A | 0623NE03 | A11-1234 | 06/2023 | 05/2026 | Nigeria |
| B | 211235 | B4-0042 | 12/2021 | 12/2024 | China |
| C | A043021 | A4-8867 | 04/2023 | 03/2026 | India |
| D | GG23004 | B4-2667P | 03/2023 | 02/2026 | India |
| E | VP16211 | A4-6297 | 04/2022 | 03/2025 | India |

The weight variation test evaluates whether the weight of tablets falls within acceptable limits, indicating uniformity in production. According to the British Pharmacopoeia (BP), the permissible weight variation range for tablets exceeding 250 mg is typically ±5% of the average weight [13,16]. Table 2 shows the weight variations of 20 tablets analysed for each brand. The average weights of the brands ranged from 267.4 mg to 953.3 mg, with Brand A exhibiting the highest percentage weight deviation, ranging from –3.85% to +4.02%. All the brands complied with the specified BP limits, as no brand exhibited more than ±5% weight variation.

**Table 2:** Weight variation of different brands of diclofenac potassium tablets (50 mg)

| Brand code | Mean Weight (mg) | Range of percentage weight variation (%) | Number of tablets outside $\pm 5\%$ of the mean weight |
| --- | --- | --- | --- |
| A | $597.0 \pm 0.015$ | -3.85 - 4.02 | 0 |
| D | $267.4 \pm 0.002$ | -1.27 - 0.97 | 0 |
| C | $886.4 \pm 0.015$ | -2.97 - 1.64 | 0 |
| D | $942.0 \pm 0.002$ | -1.16 - 1.06 | 0 |
| E | $953.3 \pm 0.014$ | -2.54 - 2.17 | 0 |

Pharmaceutical tablets must possess sufficient strength to endure mechanical stress, abrasion, chipping, and breakage during manufacturing, processing, and shipping [17]. The friability test is utilized to assess the ability of tablets to maintain their inter-particulate bonding under such mechanical stresses. The friability test conducted on five brands of diclofenac potassium tablets demonstrated that all brands were sufficiently durable to retain their structural integrity. The percentage friability ranged from 0.0213% to 0.8198% (Table 3). According to official standards, a friability test is deemed acceptable if the total weight loss after testing does not exceed 1% [16]. All tested samples of diclofenac potassium passed the friability test, achieving a 100% compliance rate.

**Table 3:** Friability test for different brands of diclofenac potassium tablets (50 mg)

| Brand code | Weight of ten tablets before friabilation (g) | Weight of ten tablets after friabilation (g) | %friability |
| --- | --- | --- | --- |
| A | 5.974 | 5.927 | 0.7867 |
| D | 2.696 | 2.694 | 0.0742 |
| C | 8.417 | 8.378 | 0.4633 |
| D | 9.393 | 9.316 | 0.8198 |
| E | 9.388 | 9.386 | 0.0213 |

While pharmaceutical tablets must possess adequate hardness to retain inter-particulate bonding and withstand mechanical stress, they should not be excessively hard or overly soft. An acceptable minimum crushing strength of 40 N is generally recommended [14]. The results of the hardness test revealed that all tested tablets required a crushing strength above this threshold. The mean crushing strengths for Brands A–E were 97.61 N, 103.50 N, 116.25 N, 109.87 N, and 103.01 N, respectively (Table 4). Although there is no universally applied strict maximum value, tablets with excessively high crushing strengths may become too hard. Tablet hardness is influenced by the amount of pressure applied during compression, which in turn impacts friability, disintegration, dissolution, and ultimately the drug’s absorption in the body [16]. Therefore, controlling tablet hardness is crucial for ensuring optimal drug performance

**Table 4:**
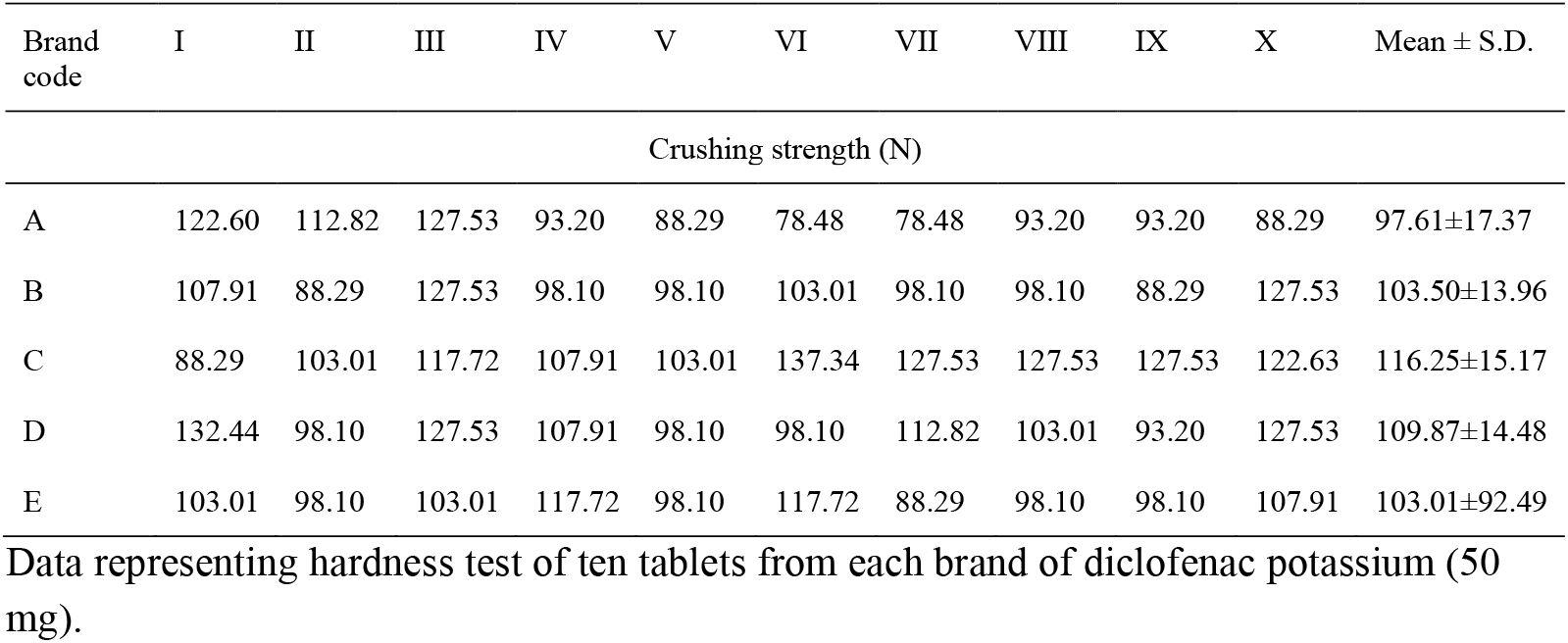
Hardness test for different brands of diclofenac potassium tablets (50 mg)

The measure of disintegration is a critical aspect of the quality control testing for tablets. The disintegration test evaluates the breakdown of tablets into smaller particles or their constituent elements within the prescribed time when placed in a liquid medium under specified experimental conditions. According to the British Pharmacopoeia (2003), uncoated tablets should disintegrate within 15 minutes [13]. For the screened diclofenac brands, the disintegration time ranged from 0.40 to 5.50 minutes. Specifically, Sample C had the highest disintegration time of 5.50 minutes, while Sample E demonstrated the lowest disintegration time of 0.40 minutes (Table 5). Several factors can influence disintegration rates, including the types of binders, disintegrants, and lubricants used during formulation [14]. Ensuring tablets disintegrate within the specified time is crucial for proper drug absorption and achieving the desired therapeutic effect. Delayed disintegration may result in therapeutic failure [14]. All tested samples met the disintegration test requirements, achieving a 100% pass rate.

**Table 5:** Disintegration test for different brands of diclofenac potassium tablets (50 mg)

| Brand code | Mean disintegration time (minutes) | Standard deviation | % Coefficient of variation |
| --- | --- | --- | --- |
| A | 3.13 | 0.72 | 23.00 |
| B | 4.42 | 1.07 | 24.21 |
| C | 5.50 | 1.20 | 21.82 |
| D | 0.40 | 0.01 | 2.50 |
| E | 0.85 | 0.50 | 58.82 |

The dissolution test is a critical quality control measure that directly impacts the absorption and bioavailability of drugs. This test determines the drug release pattern from tablets, which is essential for predicting the drug’s therapeutic performance. The percentage drug release of selected diclofenac potassium brands was calculated using the calibration curve of standard diclofenac potassium (Figure 1), which demonstrated a linear relationship with a regression coefficient R^2^=0.998. According to recommended standards, not less than 70% of the labelled amount of diclofenac potassium should be released within 45 minutes [9,13]. The dissolution pattern of each brand was observed over 60 minutes. Among the tested brands, Sample B achieved the highest drug release at 45 minutes, with 70.44% released. However, Brands A, C, D, and E failed to meet the standard, releasing only 51.2%, 52.21%, 30.53%, and 46.17%, respectively (Figure 2). Although these brands might release their drug content at later times, delayed drug release can negatively affect the onset of therapeutic action. Disintegration rate is also an important parameter for assessing bioequivalence. The presence of multiple brands in the market can create confusion among healthcare professionals and patients regarding the choice of brand and the interchangeability of products [17]. Despite all the brands having the same labelled claim, they are not bioequivalent as they do not have equal drug release profile. Only Brand B met the recommended dissolution standard, while Brands A, C, D, and E failed the test.

**Figure 1:**
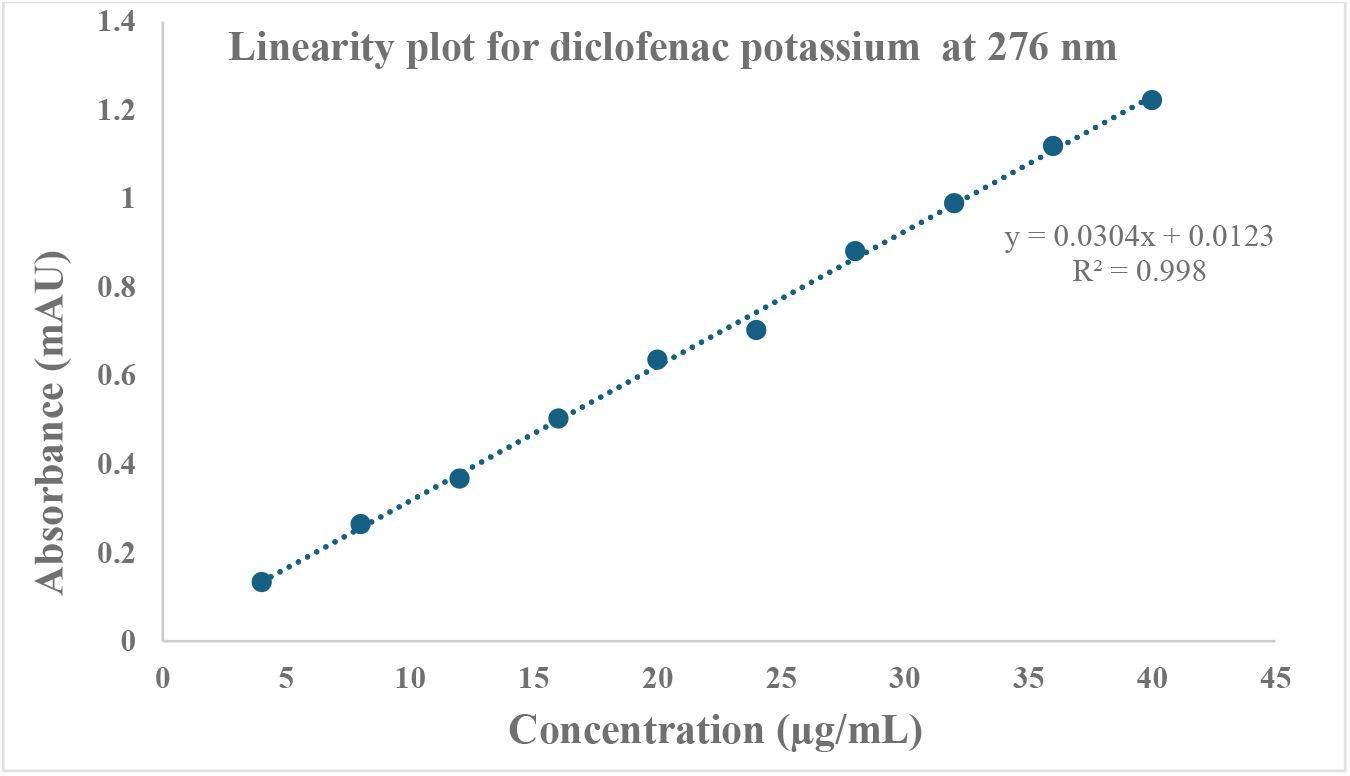
Calibration curve for standard diclofenac potassium at 276 nm showing a linear relationship between known concentrations of diclofenac potassium and absorbance using UV-Visible spectrophotometer.

**Figure 2:**
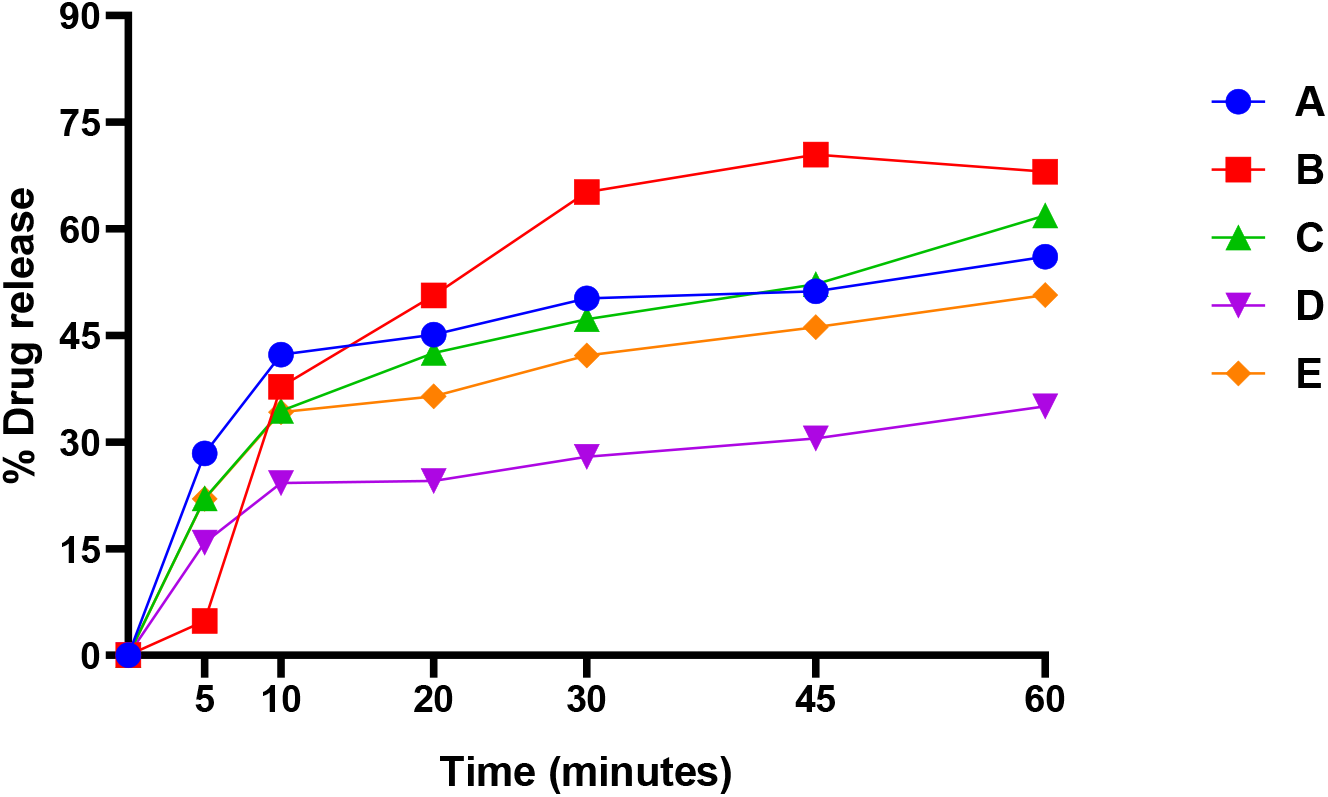
Graph of dissolution profile for different brands of diclofenac potassium (50 mg)

To evaluate the content uniformity of the selected diclofenac brands, an HPLC assay was employed. HPLC is a robust analytical technique capable of performing both qualitative and quantitative analyses of pharmaceuticals, as well as compound identification [18]. The quantification of diclofenac content for each brand was based on the calibration curve shown in Figure 3, which demonstrated a strong linear relationship between concentration and peak area, with a R^2^=0.9995. Samples A, B, C, and E successfully met the percentage purity requirements with values of 97%, 95.5%, 100%, and 98.5%, respectively (Table 6), aligning with the BP stipulated range of 95.0%–105.5% [19]. However, Sample D failed the test with a percentage content of 75% (Table 6).

**Table 6:** Content purity analysis of different brands of diclofenac potassium tablets (50 mg) using HPLC.

| Brand Code | Theoretical amount mg/ml | Experiment amount mg/ml | Label claim | Conc. (mg) | Percent content (%) | Status<br>B.P specification<br>95 % - 105% |
| --- | --- | --- | --- | --- | --- | --- |
| A | 0.2 | 0.194 | 50mg | 48.5 | 97 | Pass |
| B | 0.2 | 0.191 | 50mg | 47.5 | 95.5 | Pass |
| C | 0.2 | 0.2 | 50mg | 50 | 100 | Pass |
| D | 0.2 | 0.150 | 50mg | 37.5 | 75 | Fail |
| E | 0.2 | 0.197 | 50mg | 49.25 | 98.5 | Pass |

**Figure 3:**
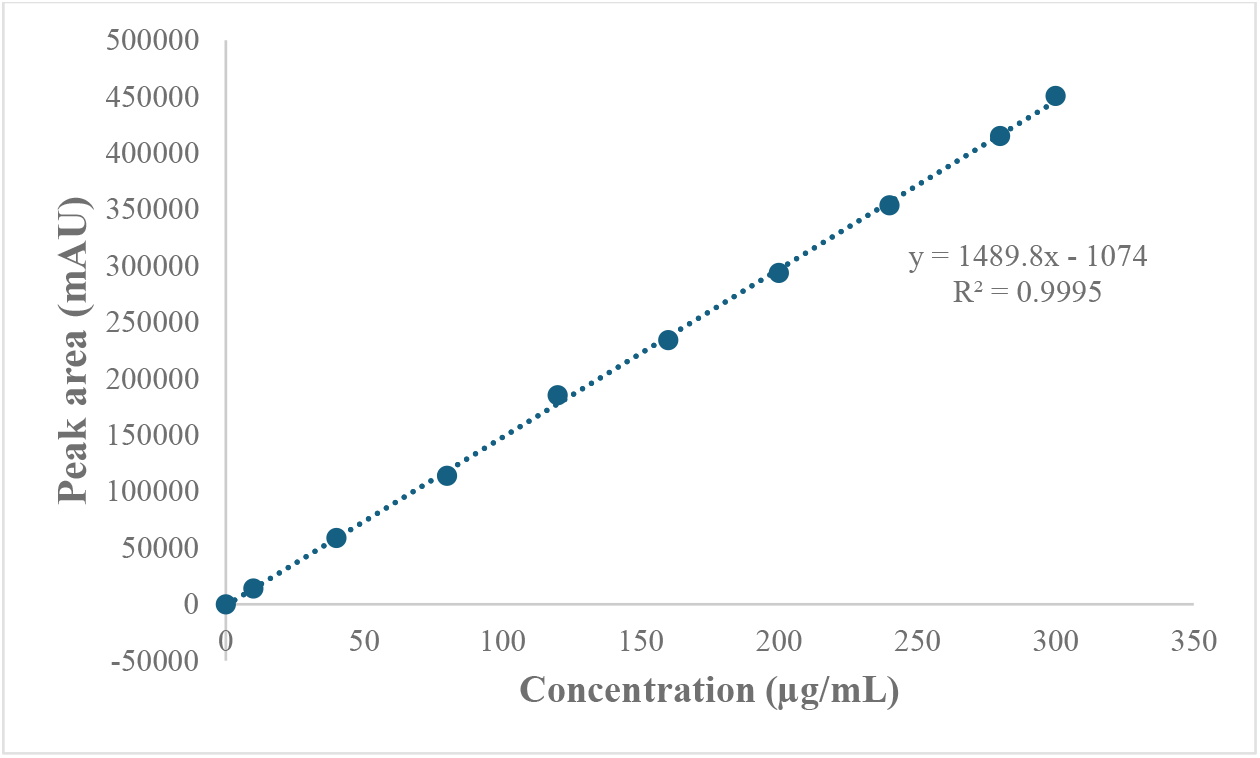
Linearity plot for standard diclofenac potassium, showing the relationship between peak areas and known concentrations for content purity analysis using HPLC.

Ensuring that pharmaceutical products contain the correct proportion of active pharmaceutical ingredients (APIs) is critical. Underdosing can lead to therapeutic failure, while overdosing increases the risk of adverse effects. Non-compliance with established quality standards may result in substandard drugs, compromising patient safety and treatment outcomes [17]. Pharmaceutical manufacturers must adhere to good manufacturing practices (GMP) and implement rigorous in-process controls to maintain product quality and ensure therapeutic efficacy.

The *in-vivo* evaluation of the analgesic activity of various brands of diclofenac potassium (50 mg) using the hot plate method revealed notable variations in the onset of analgesic effects among the treated animals. At 15 minutes post-treatment, all brands, except Brand A, exhibited a statistically significant reduction in pain compared to the negative control group (Table 7). However, no significant differences in analgesic efficacy were observed between the tested brands and the positive control at this time point. The rapid onset of analgesic action is crucial for patient relief, and this can be influenced by factors such as the tablet’s disintegration and dissolution profile, as well as physiological variations in the animals’ response to the drug [20]. By 60 minutes post-treatment, all brands demonstrated a significant reduction in pain relative to the negative control group. Notably, Brand E showed analgesic activity comparable to the positive control, while the other brands exhibited a lower degree of efficacy (Table 7).

**Table 7:** *In-vivo* analgesic activity of different brands of diclofenac potassium tablets (50 mg) on experimental rats.

| Group | Brand code | Reaction time in seconds (Mean ± S.D.) | | | | |
| --- | --- | --- | --- | --- | --- | --- |
|  |  | 0 min | 15 min | 30 min | 45 min | 60 min |
| I | A | 12.6±1.36 | 12.3±1.17 | 12.7±0.47<sup>b</sup> | 13.4±1.02<sup>b</sup> | 15.5±0.80<sup>a b</sup> |
| II | B | 12.06±0.91 | 13.2±0.75<sup>a</sup> | 14.0±0.63<sup>b</sup> | 15.3±0.76<sup>a b</sup> | 16.8±0.53<sup>a b</sup> |
| III | C | 12.02±0.45 | 13.0±0.63<sup>a</sup> | 13.5±0.5<sup>b</sup> | 14.7±0.50<sup>a b</sup> | 16.0±0.63<sup>a b</sup> |
| IV | D | 12.03±0.03 | 14.5±0.50<sup>a</sup> | 14.9±0.41<sup>a</sup> | 16.3±0.50<sup>a</sup> | 17.5±0.50<sup>a b</sup> |
| V | E | 11.93±0.04 | 13.1±0.31<sup>a</sup> | 15.2±0.35<sup>a</sup> | 16.9±1.06<sup>a</sup> | 19.2±0.75<sup>a</sup> |
| VI | Negative control | 12.03±0.06 | 11.64±0.49 | 12.86±0.75 | 11.9±0.41 | 13.3±0.50 |
| VII | Positive control | 11.34±0.03 | 13.5±0.50 | 15.7±0.49 | 17.0±0.63 | 19.7±0.76 |

Data presented as mean ± standard deviation (S.D.). Statistical significance at $p < 0.05$ , $n=5$ . <sup>a</sup>statistically significant when compared to negative control and <sup>b</sup> statistically significant when compared to positive control.

These findings reveal the importance of continuous quality control testing for pharmaceutical products. Manufacturers must adhere to stringent good manufacturing practices, while regulatory bodies play a vital role in ensuring compliance with established specifications. This is essential for guaranteeing that only safe and effective products are made available to the market.

## Conclusion

In conclusion, all the selected brands passed the uniformity of weight, friability, hardness, and disintegration tests. However, Samples A, C, D, and E failed to meet the official standard for their drug release profile. All brands, except for Brand D, passed the content purity test. Additionally, all brands demonstrated significant analgesic activity *in-vivo*, indicating their potential therapeutic efficacy. These findings emphasize the need for continuous quality control and adherence to established standards to ensure both the safety and effectiveness of pharmaceutical products.

